# Benign Taste Experience Enhances Cortical Response Reliability during Aversion Conditioning Towards a Novel Taste

**DOI:** 10.1101/2023.12.19.572413

**Authors:** Veronica L. Flores, Emma Barash, Donald B. Katz, Jian-You Lin

## Abstract

Learning depends on more than the association of paired stimuli in a vacuum. For example, benign experience with a taste stimulus weakens future conditioned taste aversions (CTA) to that taste—a phenomenon known as latent inhibition (Lubow and Moore, 1959). Our work has revealed a similar phenomenon in rodents whereby benign experience with tastes significantly enhances later CTA to a novel taste (Flores et al., 2016; Flores et al., 2018). This phenomenon suggests that taste experience broadly alters future taste processing in gustatory cortex (GC). To delve into this possibility, we previously have used *in-vivo* electrophysiology in female Long-Evans rats, recording GC activity during benign taste experience. These recordings revealed an increase in the discriminability of GC responses to both familiar and novel tastes following taste experience (Flores et al., 2022). Here, we build upon these earlier results to present a potential mechanism for this increased discriminability and salience—a reduction of cortical response variability of single-neuron responses, and enhanced coherence of ensemble responses, to the novel taste. We go on to demonstrate that the differences between pre- and post-CTA responses are greater following benign taste experience. These results suggest that benign taste experience impacts GC responses towards a novel taste pre-and post-CTA conditioning by increasing taste coding reliability at both the single-neuron and ensemble levels.

**Significance Statement:** Animals and humans readily learn the consequences of consuming a specific taste and react by changing their behaviors. However, those changes often vary and can be influenced by previous experience. For example, our previous work has shown that inconsequential taste experiences increase the probability of a strong taste aversion. The work presented here evaluates how benign taste experience can impact cortical processing dynamics using *in-vivo* electrophysiology in freely behaving rats. We report that seemingly inconsequential taste experience impacts cortical plasticity. This unravels a new area of chemosensory research and may shed light on what differentiates the neural dynamics of strong and weak taste aversion learning.

## Introduction

Behavioral modifications in response to experience play an essential role in animal survival and decision making. Researchers studying these modifications typically focus on experiences linked to specific associated consequences, such as conditioned taste aversion (CTA), wherein an animal learns to avoid a particular taste (the conditioned stimulus; CS) that was previously associated with malaise (unconditioned stimulus or US; Garcia et al., 1955). The associability of the conditioned stimulus in CTA learning is nearly unique in that a single paired experience with malaise causes robust and reliable learning.

But this association process does not exist in a vacuum—rather, it itself is influenced by prior experience. Simple inconsequential experiences with the eventual CS, for instance, weakens its associability (a phenomenon known as latent inhibition; Lubow and Moore, 1959; Lubow, 1973; De la Casa and Lubow, 1995). More recently, we have described a complementary effect, wherein experience with salty and/or sour tastes increases the associability of novel sucrose (leading to an enhanced CTA in a phenomenon that we will refer to as Latent Enhancement of CTA or LE).

Relative to latent inhibition (Naor and Dudai, 1996; Miranda et al., 2003; Reilly and Schachtman, 2009; Yiannakas and Rosenblum, 2017) what we know about the neural basis of LE is minimal. It is likely that the effect involves Gustatory cortex (GC), which is a central part of the taste learning and behavior circuit (Yamamoto et al., 1995; Berman and Dudai, 2001; Uematsu et al., 2015; Levitan et al., 2016; Li et al., 2016; Wiaderkiewicz and Reilly, 2023). Consistent with this prediction, Flores et al. (2018) reported elevated levels of CTA-related c-FOS expression within GC when animals were exposed to benign tastes prior to learning, and showed that optogenetic inhibition of GC activity during benign taste experience abolished LE. Furthermore, GC spiking responses to presentations of novel stimuli change as a result of benign taste pre-exposure. GC taste responses normally evolve through various coherent firing epochs related to the detection, identity, and palatability of tastes (Katz et al., 2001, 2002; Jones et al., 2007); following taste experience, activity within the palatability epoch for both novel tastes and each of the experienced tastes becomes more distinct (e.g., with experience, water became more discriminable from salty tastes which was more discriminable from sour tastes; Flores et al., 2022).

Discriminability is, of course, a measure comparing multiple and different taste responses and thus one factor we were able to study during benign taste experience. Discriminability is not an appropriate variable when trying to understand changes towards a single stimulus (such as the impact of taste experience on an aversion towards a single novel taste). Given that the aversions are typically stronger after taste experience, our research questions here seek to link the increase in experience-induced discriminability seen prior to aversion learning to potential mechanisms that can help shed light on why stronger learning occurs following taste experience. Our results reveal that benign taste experience not only increases the discriminability of pre- and post-CTA sucrose responses, but it also increases the reliability of GC responses towards novel sucrose pre-CTA. This increase in reliability is evident through decreased trial-to-trial variability within single-neuron responses and increased neuron-to-neuron coherence of ensemble activity. Furthermore, despite a relatively compressed range of learning strengths (a result of small behavioral sample and optimization of methods for electrophysiology), this increase in reliability correlated with stronger learning. These results show that benign taste experience impacts GC response plasticity towards a novel taste pre-and post-CTA learning in a way that can may offer insight into both the increase distinctiveness of responses and the LE phenomenon. Future research is discussed addressing whether this increased reliability in GC taste processing is explicitly linked to LE.

## Methods

### Subjects

Adult female Long Evans rats (N = 8; 250-315g at time of surgery) from Charles River Laboratories served as subjects for these experiments. On their arrival, the animals were housed in a temperature-controlled vivarium with 12:12 light-dark cycle (light on at 8:00). Animals had *ad libitum* access to food and water prior to all experiments, during which water consumption was restricted by giving 15 mL of water each day following experimentation. All procedures were conducted in accordance with the guidelines established by the Brandeis University Institutional Animal Care and Use Committee.

### Taste Stimuli

Taste solutions consisted of 0.1 M sodium chloride (NaCl; salty taste), 0.2 M citric acid (sour taste), 0.2 M sucrose (sweet taste used for CTA learning/testing), and distilled water (water). We chose these concentrations to remain consistent with our previous investigations of benign taste experience (Flores et al., 2016; Flores et al., 2018; Flores et al., 2022).

### Experimental Design

The experimental paradigm is five days long and is summarized in Figure 1. On each day, experiments were conducted between 12pm and 2pm in a Plexiglass experimental chamber separate from the rats’ home cages. During the experimentation, rats were water deprived for 19 hours/day to ensure task engagement. *In-vivo* electrophysiological recordings were collected during each of the five days of the experimental protocol.

**Figure 1.**
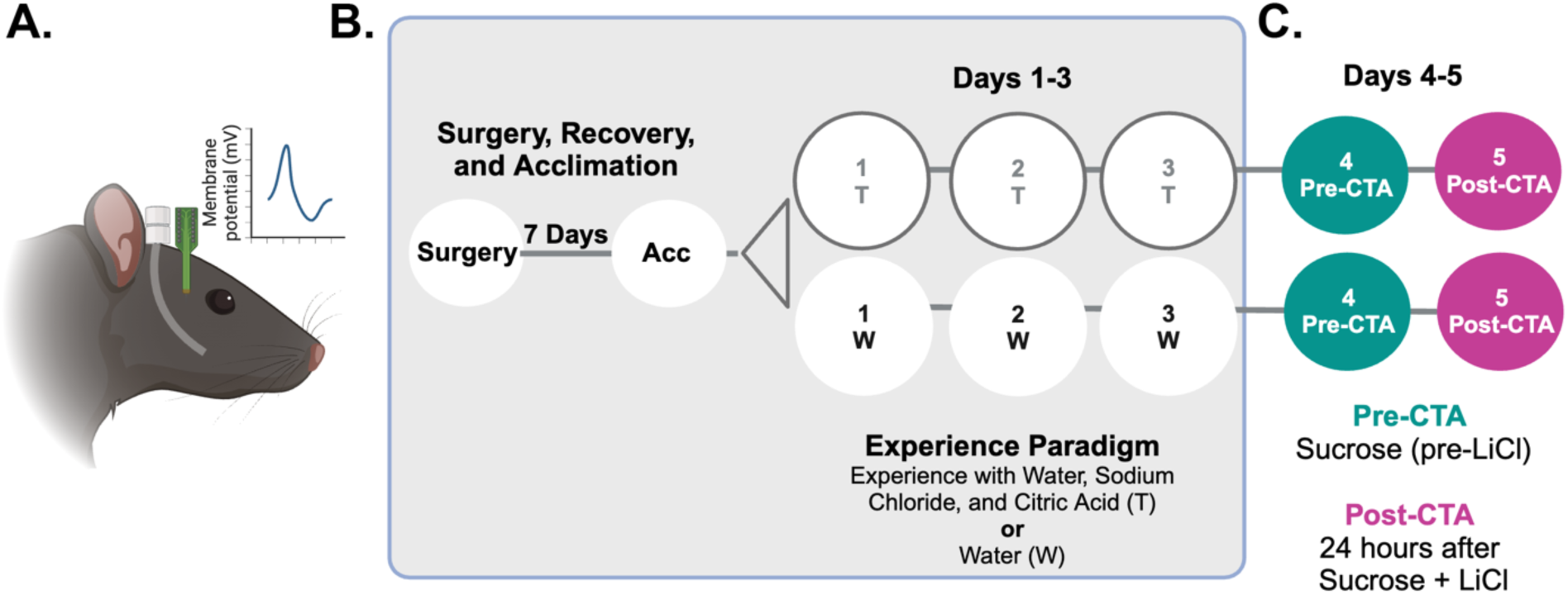
Electrophysiology Paradigm. A. Representation of Long Evans rodent with intra oral cannula and electrode implantation. B. Focus of analyses in Flores et al., 2022. Schematic of the experimental timeline occurring before the pairing of sucrose and lithium chloride: implantation surgery, 7 days of recovery, 1 day of acclimation (Acc), 3 days of experience with water (W) or tastes (T: water, sodium chloride and citric acid). C. The focus of this manuscript is on neural recordings taken during novel sucrose consumption (pre-CTA) and one day of sucrose consumption following the pairing of sucrose and lithium chloride (post-CTA).

#### Taste Experience Sessions

Extensive details regarding taste experience sessions can be found in Flores et al. (2022). In brief, after acclimation to IOC delivery via water presentation (Acc in Figure 1B), rats were given trials of either water (control group herein referred to as the water-experienced group) or a taste battery (water, sodium chloride, and citric acid: herein referred to as the taste-experienced group) for three consecutive days (Days 1 – 3, Figure 1B). Each session included 120 deliveries of water or the taste battery (40 deliveries of each water, sodium chloride, and citric acid); each delivery involved 40ul bolus of fluid, separated by 15-s inter-trial intervals. Rinse trials were not used as this length of interval is sufficient for responses to return to baseline in the active tasting, non-anesthetized paradigm (Katz et al., 2001; Jones et al., 2007; Moran and Katz, 2014).

#### Pre-CTA Session

Twenty-four hours after the 3^rd^ taste experience session, rats were given novel sucrose (pre-CTA, Figure 1C). Rats were first presented with 5-min access to a bottle of sucrose in the home cage (maximum consumption was 15ml), after which they were moved to the experimental chamber to receive 120 trials of sucrose via IOC (delivery schedule identical to experience sessions); the bottle consumption ensured the development of CTA, while the IOC deliveries were best for simultaneous *in-vivo* recording of GC activity. Thereafter, each rat received a subcutaneous injection of lithium chloride (LiCl, 0.3 M, 0.5% of bodyweight) to induce CTA learning before being returned to their home cages.

#### Post-CTA Session

On the fifth day (post-CTA, Figure 1C), rats were again given access to 15 ml of sucrose for 5 minutes in the home cage. Immediately following behavioral testing animals were placed in the experimental chamber to receive 120 trials of sucrose via IOC while again recording GC taste responses. Of the seven rats who were used to collect electrophysiological data from, five were included in the behavioral analysis to measure aversion learning (one rat was removed due to being a statistical outlier and one rat was removed due to an experimental error where pre-and post-measurements were not acquired).

### Acquisition of In-Vivo Electrophysiological Data

Electrophysiology recordings were recorded from each animal during the five days of the experimental paradigm (see details below) within a customized Faraday cage (8.5 x 9.5 x 11.5 in), located in an isolated room adjacent to the animal holding space. The Faraday cage was equipped with one desktop computer (PC) and one Raspberry Pi computer (Model 3B). The Pi computer controlled the taste stimulus presentation by manipulating the opening time and duration of solenoid valves for each taste delivery. The PC collected all electrophysiological recordings from the headstages attached to implanted electrodes (Intan Technologies – RHD2132, see implantation details below). These headstages were capable of sampling every electrode at 30 kHz and digitizing the signal before saving it to the PC. Subsequently, the collected recordings were sorted and analyzed offline using a set of Python analysis codes developed according to the current experimental design (for more details, refer to Mukherjee, 2017).

Only neurons meeting certain criteria were saved in hierarchical data format (HDF) for subsequent data analysis to ensure that only stable and isolated neurons were used. The criteria set the signal-to-noise ratio to higher than 3:1 and required spiking activity to exhibit a clear refractory period of 1 millisecond. We then compared spike times and removed neurons that exhibited significant overlap (at least 50% overlap). This cautious approach minimized the possibility of including the same neurons detected by more than one electrode channel.

### Analysis of Electrophysiology Data

After histological confirmation of electrode placement, one animal was removed from the electrophysiological data set leaving N = 7 rats used to collect electrophysiological data from N = 261 neurons.

In Flores et al. (2022) we examined cortical responses during the three days of benign taste experience. Here, we focus on days four and five in the same rodents: the *in-vivo* recordings of gustatory cortex towards novel sucrose pre- and post-CTA learning that occurred after benign taste experience with salty and sour tastes compared to animals who experienced water alone. Similar to those in the companion paper, single neuron data was subjected to 3 steps of previously described analyses (pre-processing, processing and post-processing) designed to characterize neural responses to stimuli (see Mukherjee, 2017 for in-depth details).

#### Single neuron trial-to-trial response reliability during CTA conditioning

For single-neuron variability, we used Fano factor (FF) to quantify the variance in taste responses across sucrose trials. As depicted in the following equation, FF is the ratio of the variance and the mean of the firing rates for 120 sucrose trials during CTA conditioning.

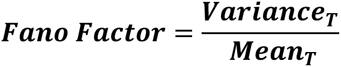

Where *Variance* and *Mean* represent the variance and the mean of firing rates across 120 sucrose trials (*T*) during the CTA conditioning session.

#### Ensemble (neuron-to-neuron) response reliability during CTA conditioning

As demonstrated in several independent works, GC taste response is dynamic (Katz et al., 2001; Jones et al., 2007; Sadacca et al., 2012) and the quality of taste coding depends on how the epochal response evolves through time. These epoch transitions are characterized with sudden, coherent changes in neuronal firing rates. The transitions can be detected in single trials and inferred from activity of an ensemble with as small as 6 simultaneously recorded neurons (Jones et al., 2007). While many statistical algorithms have been applied to detect state transitions in neural spiking data (hidden Markov Modeling, Jones et al., 2007), here we used Bayesian Changepoint Analysis (BCA) to determine the presence of transitions in ensemble taste responses. Unlike many other methods that provide a single point estimate through maximum likelihood estimation, we chose BCA because, as demonstrated below, it offers a distribution of parameter estimates. This distribution effectively captures how certain the model is about the parameter estimates (in our case, transition times) – the wider the distribution, the more uncertain the model is about the estimates which is crucial for accurately characterizing the quality of state transitions.

BCA is written in the probabilistic programming language pymc3 (Salvatier et al., 2016). The pipeline of implementation of this method is summarized in Figure 2A: taste-evoked spike trains were collapsed into consecutive 50ms bins for each trial and GC neuron, and a Poisson likelihood estimate was used to model the binned spike counts. Similar to a hidden Markov model, the changepoint models consisted of two sets of latent variables: (1) the emission variables (i.e., firing rates) and (2) the changepoint position variables (i.e., in which time bin changepoints occur); equations and the dependencies between these variables are detailed in Figure 2B. Briefly, the mean firing rates for each neuron is modeled independently; emissions for neurons in each state are dependent on neurons. As such, these variables are hierarchically organized to better constrain the space of emission values during different states for each neuron (i.e., the model knows the emission for State 1 for Neuron 1 will be similar to State 2 for the same neuron). The changepoint positions are defined assuming that the distribution of hyperparameters (τ_a_, τ_b_) which are shared across all trials for a given changepoint. Finally, the latent emissions and changepoints are combined to generate time series of firing rates with sequential states (f(λ,τ)) is a deterministic function, which is used to evaluate the likelihood of the data (obs) given the latent variable. The output of the model fit is posterior distributions of the parameters. Herein we use the mean of the distribution as the estimate of transition time, while the variance of the distribution (termed highest density interval [HDI]) serves as the estimate for the quality of the transitions. Figure 2C shows the fitting results from 10 consecutive sucrose trials of a single recording session. A modularized pipeline was used to fit and analyze the models across datasets (https://github.com/abuzarmahmood/pytau/tree/development/pytau), enabling efficient comparisons between rats with and without prior taste experience.

**Figure 2.**
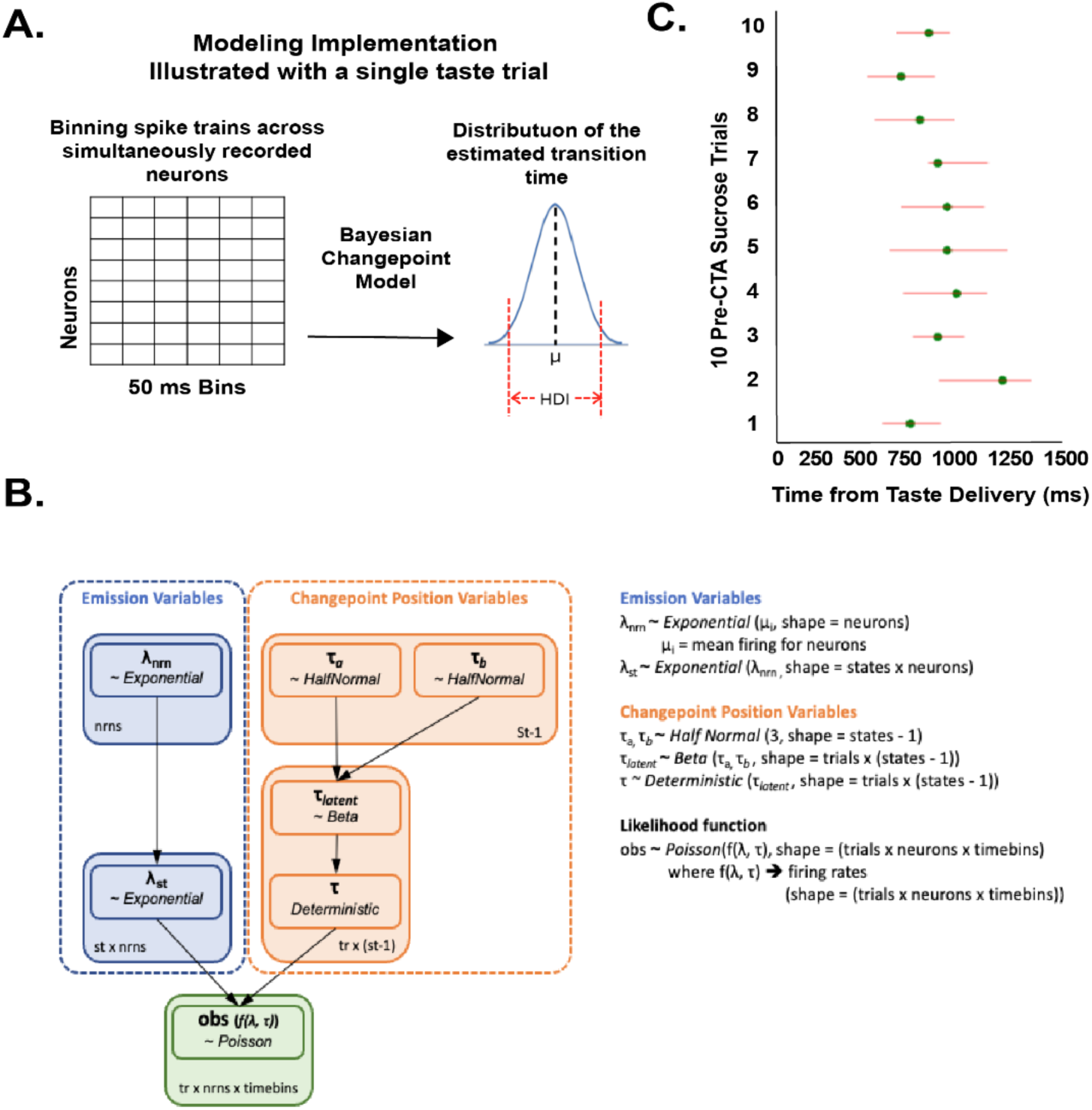
Model of Bayesian Changepoint Analysis (BCA) and its Implementation. A. Schematic of the model execution pipeline. Data (taste-evoked spike trains) were first collapsed into 50-ms bins (summation of the spikes in each bin) for each trial and each neuron before feeding into the BCA. The result of the fitting process are distributions of parameter estimates, from which we can gather estimates of when state transitions are most likely to occur (indicated by mean of the distribution [μ]), and how certain the model is about the estimate (indicated by an interval containing 95% of the values of the posterior distribution, termed highest density interval [HDI]). B. Dependencies between the random variables for constructing the BCA model are shown within a schematic: (number of neurons – nrns), (number of states – st) and (number of trials – tr). C. The estimates of transition times (green dots) and HDI ranges (red horizontal lines) for 10 consecutive trials of a recording session.

#### Evaluating single-neuron stability across days

To determine if waveforms recorded from a single electrode represented the same neuron across 24 hours (from pre- to post-CTA sessions), we performed a previously described waveform-based held-unit analysis (of conditioned taste aversion; Grossman et al., 2008; Moran and Katz, 2014). A principal component analysis was used to map the shape of each waveform in 2D such that each waveform formed a “cluster”. Clusters that overlapped over the two sessions indicate a high potential of similarity (suggesting that they are the same neuron). Separation of each cluster was calculated using a non-parametric clustering statistic (*J*_3_) for the first and last third time periods of activity for each waveform during each of the sessions (2) being compared. Waveforms were considered “held” across sessions if the *J*_3_ value between sessions was less than 90% of the entire *J*_3_ distribution.).

After identifying held-units across sessions, we examined whether experience impacts the way that single unit GC taste-responses change pre- and post-CTA learning, we measured the Euclidean distance (ED) between each single neuron’s taste response pre-CTA and the average firing rate response (first 2 seconds of the taste response) across all neurons (templates) for each taste delivery trial of sucrose post-CTA. Measuring Euclidean Distance helps determine the similarity between two same-length arrays (herein referred to as [post-CTA array] and [pre-CTA array]); for our purposes this measurement helps us determine how CTA learning impacts responses towards sucrose in single neurons. The larger the distance, the greater the difference between neural responses towards sucrose pre- and post-aversion learning. Here, the array lengths are the number of single neurons held across the 24-hour period (taste experience: N =11 and water experience: N=10. SPSS was used to perform one-way ANOVAs comparing Euclidean distances between sucrose responses pre- and post-CTA.

## Results

Here, we ask whether we can identify an aspect of responding to individual tastes that is: 1) changed by pre-exposure to NaCl/CA; 2) non-comparative; 3) good explanation for discriminability enhancements; and 4) potentially related to strength of learning.

We have previously shown that benign taste experience enhances the discriminability of GC responses to both familiar and novel tastes (Flores et al., 2022). Here, we sought to identify an aspect of GC response that could account for the enhancement of taste discriminability and explain the strength of CTA learning. As shown below, the data presented here implicates response reliability as this aspect. We report results based on the analysis on spiking activity from 261 neurons total; 131 neurons pre-CTA (38 from the water experience group and 93 from the taste experience group) and 130 neurons post-CTA (30 from the water experience group and 100 from the taste experience group).

### The Impact of CTA Learning on GC Taste Responses Becomes Stable Following Taste Experience

The sample size for this study was designed to power the analysis of neural activity rather than behavioral measures of learning, therefore the fact that we did not see a statistically significant stronger aversion in the taste experience group is expected (although the aversion was notably stronger following taste experience, *t* (3) = -2.328, *p* = 0.102, taste experience post CTA consumption mean = 0.39mL and the water experience post CTA consumption mean = 1.00mL). Although this finding was inconsequential to our current research questions, it remains important, nonetheless, to provide at least basic replication of previous results, showing that taste experience enhances coding distinctiveness (Flores et al., 2022). To avoid an artifact of sampling from different neural populations, we specifically compare the impact of CTA conditioning on GC neuronal firing between taste experience groups but only on the neurons that were held across the 24 hours between pre- and post-CTA sessions. Figure 3A shows evidence that aversions were acquired occurred (collapsed across all rodents (N=5, see methods under *Post-CTA Session*) allowing us to then reliability examine differences between pre- and post-CTA responses. An Independent Samples *t*-test with a 95% confidence interval for the mean difference confirmed that sucrose consumption post-CTA was significantly lower than sucrose consumption pre-CTA ((*t* (8) = 10.29, *p* < 0.001) across all rodents.

**Figure 3.**
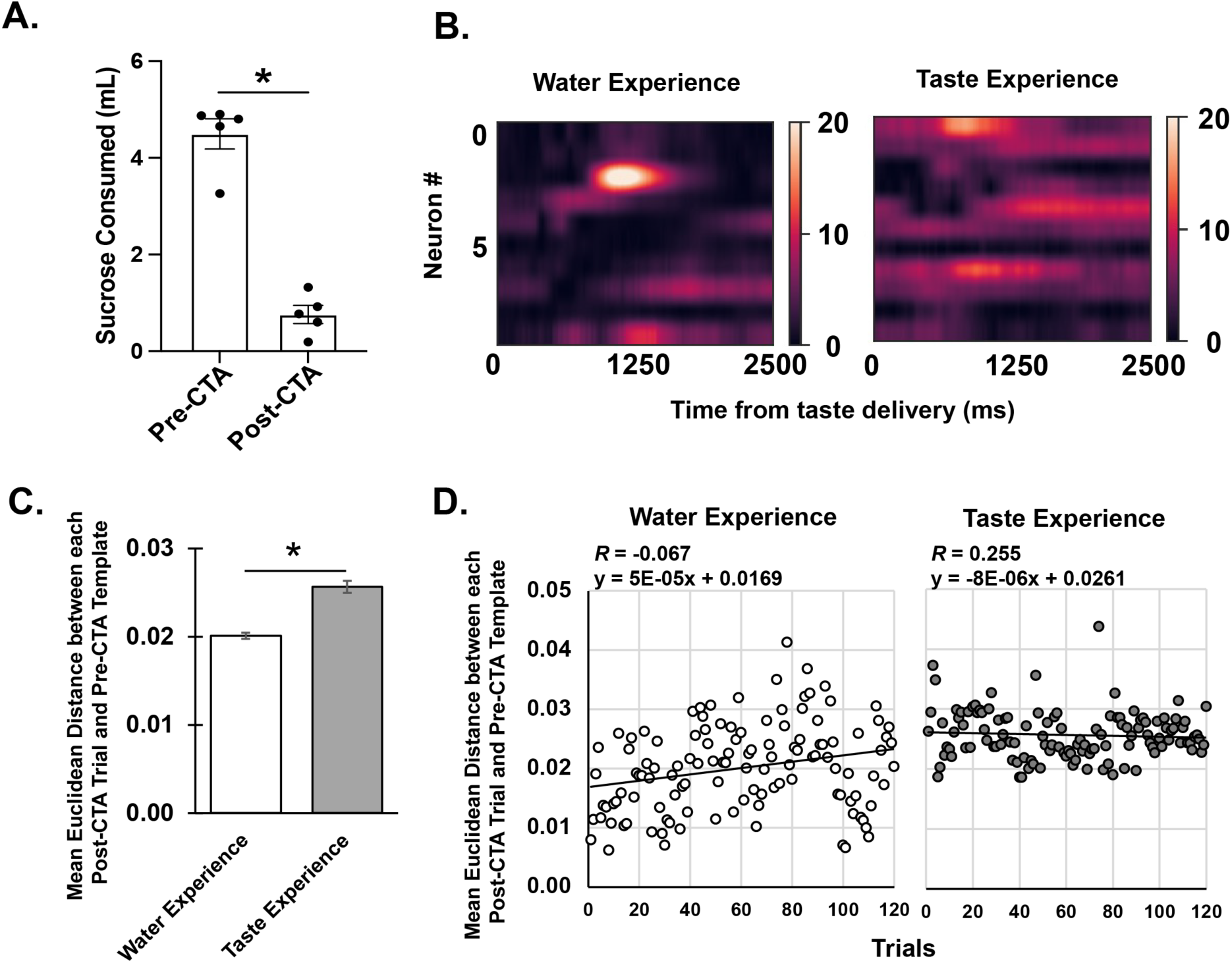
Taste Experience Stabilizes Single-Neuron Responses. A. Demonstration of CTA strength from recorded animals. Bar graph demonstrating raw sucrose bottle consumption pre-and post-CTA (n=5). Consumption of sucrose post-CTA was significantly less than sucrose consumption pre-CTA across all animals. B. Heat maps demonstrating the absolute change in firing rate for all held units for all trials. C. Bar Graph demonstrating changes in single-neuron responses to sucrose pre- and post-CTA in both water and taste experience groups. Bars signify mean Euclidean distance between post-CTA sucrose responses and a single pre-CTA template per group. Changes are between water (n = 10) and taste-experienced groups (n = 11). D. Scatterplots with lines of best fit demonstrating the Euclidean distance between each post-CTA trial and the pre-CTA template for water experience (left) and taste experience (right). Error bars represent standard error of the mean. * *p* < .05.

Even in this small sample of neurons (n = 11 taste experience, n = 10 water experience) taste experience did prove to significantly enhance response plasticity: a heatmap showing learning-related changes in firing rates across all neurons held across conditioning clearly suggests that neurons changed more with taste experience (Figure 3B); inspection of the figure suggests that CTA-driven response changes are larger and of longer durations, indicated by brighter and warmer colors in the heatmap, for taste experienced rats than those in taste naïve subjects. These differences are summarized in Figure 3C, which plots the Euclidean distances (ED; with firing rates of each neuron as the dimensions) between single-trial post-CTA sucrose responses and pre-CTA sucrose responses (averaged across trials) for both taste- and water-experienced groups (for details see Methods). An independent samples t-test performed on these data reveals that post-CTA responses are more distinct (greater ED) from pre-CTA responses for neurons from rats with taste experience than for neurons from water-experienced rats (t (238) = 7.25, p < 0.001). Taste experience was effective and powerful.

Investigation of these changes also revealed evidence that responses in taste-experienced rats were more stable than in water experienced rats. Figure 3D shows EDs between each post-CTA trial and each pre-CTA response for both experience groups. Visual inspection of this figures shows that EDs gradually increase across trials for rats in the water experience group (*r* = 0.255, *p* = 0.005); this across-trial change, however, was not observed in the taste experience group (*r* (120) = -0.067, *p* = 0.469). This pattern of results suggests benign taste experience leads to more reliable GC responses towards a novel taste involved in aversion conditioning.

### Gustatory Response Reliability Towards a Novel Taste Increases with Taste Experience

Of course, these results were observed in a small sample of neurons held across the multiple days of the learning process. It’s also worth noting that the observed differences in GC taste response reliability could be interpreted in multiple ways. We therefore decided to perform an extensive set of analyses on the entire sample of neurons (n = 261) to better characterize how taste experience alters response reliability.

At the single neurons level, we investigated whether trial-to-trial variability in sucrose-evoked responses are reduced after benign taste experience. In this case, the reliability of GC responses was quantified using Fano factor (FF; (Shadlen and Newsome, 1994, 1995, 1998), which measures firing rate variability, across 120 sucrose trials during the pre-CTA session (see Methods for formula).

Between-group comparisons of Fano Factors were analyzed across each 400ms time bin of the taste response to capture specific changes by epoch (Figure 4A). A mixed 2-way ANOVA revealed a significant main effect of experience (*F* (1, 129) = 6.43, *p* = 0.012) and Time bin (*F* (5, 645) = 1.96, *p* = 0.082), as well as a significant interaction of Group and Time (*F* (5, 645) = 3.35, *p* = 0.005). Post-hoc comparisons with Bonferroni corrections further revealed that the water-experienced group has significantly larger FFs than the taste-experienced group at baseline (-400-0 ms; *p* < 0.05); this difference then collapses at the beginning of the responses (a result consistent with prior meta-analysis of a range of preparations; Churchland et al., 2010), only to quickly return, becoming significant again by the time of transition to the late palatability epoch (800 – 2000 ms; *p* < 0.05). These results reveal that responses to sucrose are more reliable after taste experience than after water experience, and that in fact taste experience changes the nature of even spontaneous GC network activity to be more stable.

**Figure 4.**
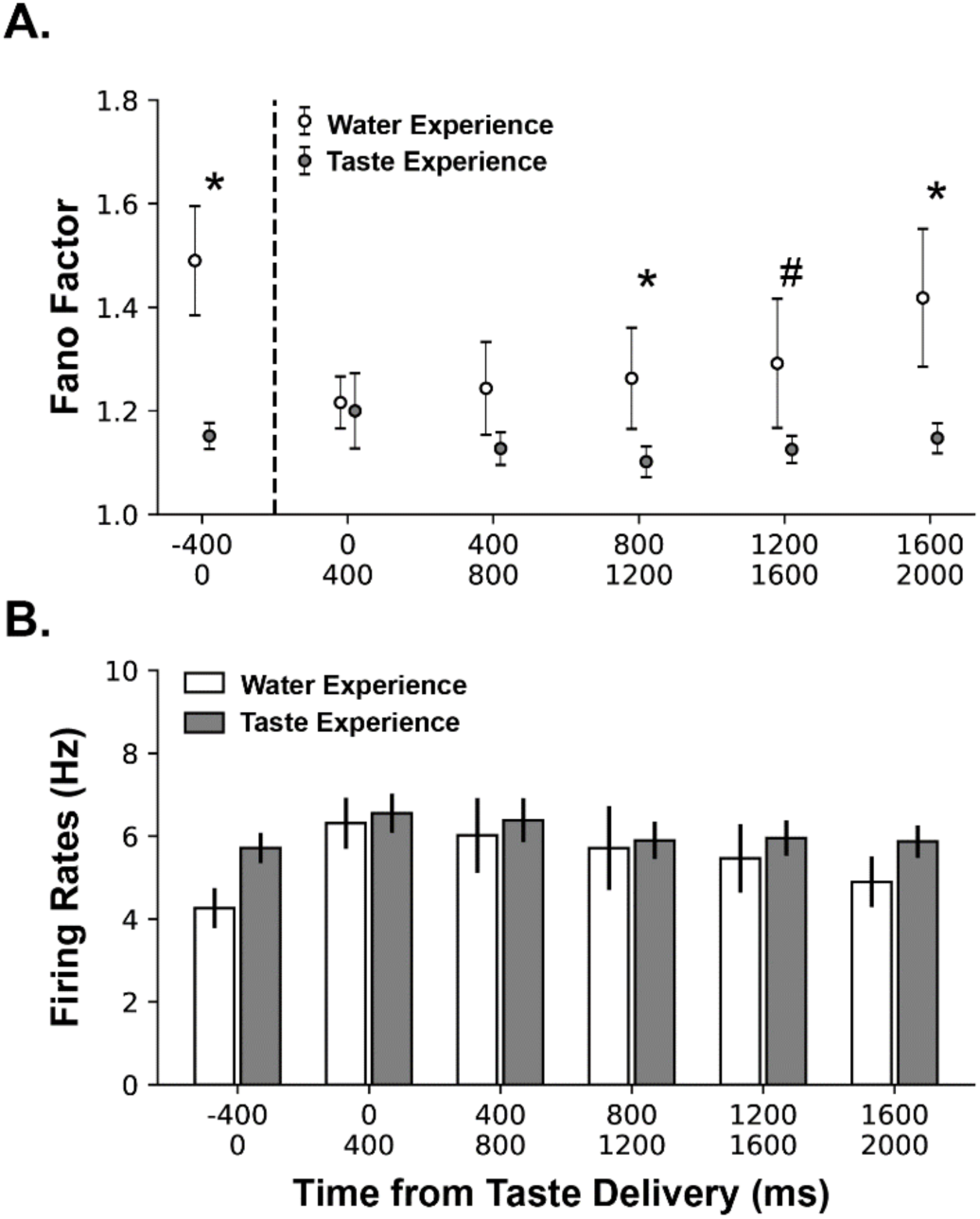
Taste experience Improves the Reliability of Sucrose Responses before CTA Conditioning. A. Mean Fano factors across each 400ms time bins between groups with either water experience (n = 38) or taste experience (n = 97). B. Mean firing rates across each 400ms time bins between groups with either water experience (n = 38) or taste experience (n = 38). * *p* < 0.05; # *p* = 0.06. Error bars represent standard error of the mean.

Fano factors are calculated as the ratio between variability and mean firing rates, which means that the observed reduction with experience could conceivably reflect changes in either of these firing properties. To test if low FFs in the taste-experienced group were in fact a result of increased firing rate instead of a reduction in variability, a two-way ANOVA was conducted to analyze the relationship between experience and firing rates. Experience with salty and sour tastes did not prove to change these firing rates in a statistically significant manner (*F* (5, 645) = 1.15, *p* = 0.33; Figure 4B). These results suggest that the reduction in FF caused by taste experience specifically reflects decreases in the variability of taste response during the palatability epoch.

In addition to the variability across trials at the single neuron level, another way that the reliability of GC taste responses can be interpreted is in terms of population (neuron-to-neuron) response dynamics within single trials (Moran and Katz, 2014; Mukherjee et al., 2019). The transitions between epochs (i.e., detection, identity, palatability) have been shown to involve ensembles of neurons changing their firing rates coherently: the more reliable the activity of the neurons in the ensemble, the more coherent the transition (see Jones et al., 2007; Sadacca et al., 2016). Increased response reliability would in this context mean a smoother and more coherent state transition within the entire ensemble. To determine whether taste experience causes such increases, we examined the impact of benign taste experience on state transitions between identity and palatability epochs pre-CTA. We targeted this transition on the basis of the previous analysis which specifically showed that the reduced variability in taste responses becomes significant in the palatability epoch, and on the basis of previous literature implicating this state transition in behavioral changes (Grossman et al., 2008; Moran and Katz, 2014; Arieli et al., 2022).

To identify this transition, we used a Bayesian changepoint analysis (BCA) with a 2-state model on the response period between 200ms to 1500ms—the period with the highest likelihood of GC activity transitioning from the identity epoch into the palatability epoch ((Figure 4A; also see Jones et al., 2007; Sadacca et al., 2016). Rather than generating a single highest likelihood of transition time estimate (which is the output of the more oft-used Hidden Markov Model; Abeles et al., 1995; Jones et al., 2007; Moran and Katz, 2014), the BCA model generates a distribution of parameter estimates for state transition time likelihood. The variance of these distributions (highest density interval, HDI) reflects how confident the model is of the transition time estimate, such that we can use HDI as the proxy for the neuron-to-neuron reliability of the transition. That is, a more reliable coherent transition is expected to result in a narrower distribution (smaller HDI) because the model is more certain about the estimation.

Figure 5B shows a representative GC neuron in which firing rates suddenly increase following an ensemble state transition (red dots in the raster plots; top panels) detected by BCA.

**Figure 5.**
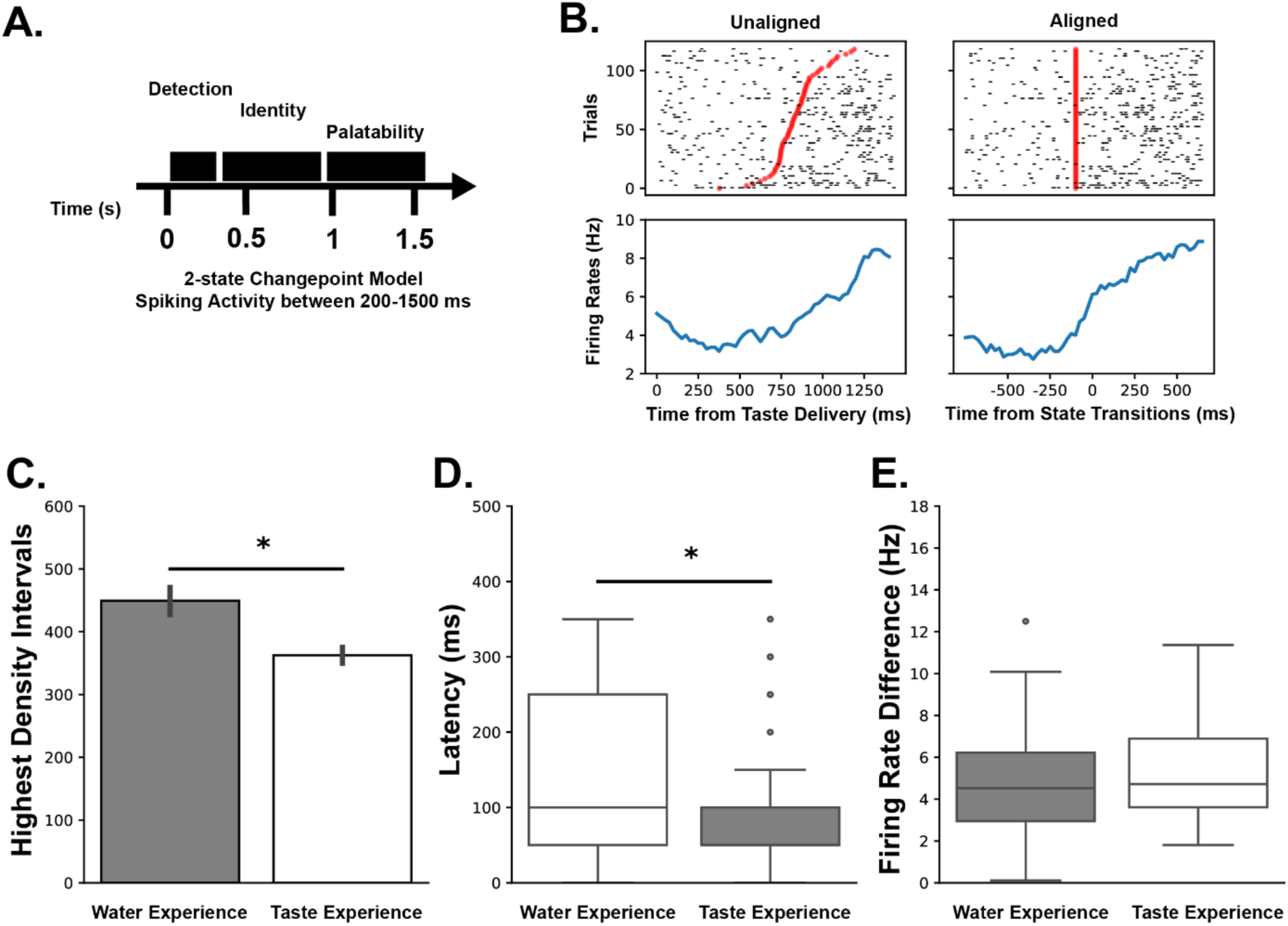
Taste Experience Sharpens GC State Transitions into the Palatability State Pre-CTA Conditioning. A. Schematic of GC response dynamic depicting that a 2-state changepoint model was used to fit spiking activity between the identity and palatability states. B. Raster plot (top panels) and the evoked firing rates (bottom panels) from a representative GC neuron. This neuron shows a sudden increase in its firing rate at different latencies across trials (Unaligned). The transition time on each trial (red dots in the raster plots) was inferred by Bayesian changepoint analysis on the activity of the entire simultaneously recorded ensemble. Once the spikes were aligned to the transition time (Aligned), the sharp change in neural activity can be better appreciated via the peri-transition histogram. C. Mean highest density intervals (i.e., the uncertainty of the model estimates of the transition time) across Water (n = 38) and Taste Experience (n = 93) Groups. D. Mean latency between single neuron transition time and ensemble transition time. E. Mean firing-rate changes across transition times across groups. * *p* < 0.05 indicating the significant between-group differences. Error bars represent standard error of the mean.

Once the spikes were aligned to the transition time (Aligned), the sharpness of the change in neural activity can be better appreciated (compare the peri-transition histogram to the right to the peri-stimulus histogram to the left). Figure 5C shows the results of the reliability analysis across our entire sample of neural ensembles (n = 6), in which the mean HDIs were plotted for water- and taste-experienced groups: neurons with taste experience have significantly smaller HDIs than those with only water experience (*t* (718) = 6.55, *p* = 0.000).

We moved on to investigating the two (non-mutually exclusive) mechanisms by which taste experience might improve state transitions (Lin et al., 2021): 1) the neurons in GC could fire in a more synchronized pattern in response to taste stimuli, such that each neuron changes its firing rate closer to the estimated time of the overall ensemble state transition; or 2) neural responses could change more drastically across transitions—the magnitude of the firing rates might change more. Only the first option would imply an actual change in neuron-to-neuron response reliability.

To determine which of these two response parameters are changed by taste experience, we ran separate changepoint models to estimate the time of state transitions in each individual neuron’s spiking activity, and then calculated the difference between each neuron’s estimated transition time and the estimated time of ensemble transition. As shown in Figure 5D, we found significantly smaller latencies between estimates from single neurons and ensembles for taste-experienced rats than water-experienced rats (*t* (67) = 2.41, *p* = 0.019). Meanwhile, when we compared the firing rates in the 200ms before and after the transitions (i.e., second possible mechanism), we observed no significant differences between rats with and without taste experience (Figure 5E; *t* (67) = -1.32, *p* =0.19). The sharpening of transitions caused by taste experience reflects an actual increase in response reliability from neuron to neuron, and not changes in the magnitude of firing-rate changes across the transitions.

### Aversion Strength Correlates with Measures of Reliability

The fact that taste experience reduces trial-to-trial variability in GC sucrose responses both offers an explanation of previous results (Flores et al., 2022) showing increased discriminability of those responses and motivates a hypothesis that high response reliability could specifically drive strong CTA learning (see Discussion).

To test this hypothesis, we evaluated the Pearson correlation between the reliability index (i.e., the Fano Factor measured during the palatability epoch) and the strength of CTA (post-CTA consumption / (pre-CTA consumption + post-CTA consumption)), without regard for group (water-experience n = 3, taste-experience n = 2; one taste-experience animal was excluded because an error led to loss of its behavioral data, and one water-experience animal was excluded for having too few neurons). Even with so few subjects, the correlation between these variables was significant (*r* = 0.82, *p* = 0.04), indicating that the strength of CTA learning is indeed a function of the trial-to-trial reliability of CS processing. Thus, the enhancement of reliability caused by experience could well be the mechanism of LE, whereby rats who receive experience to NaCl and citric acid learn stronger CTAs (i.e., drink less) than those who only experience water. Future research should further examine this prediction.

## Discussion

This research reveals that taste experience impacts the cortical processing of novel taste learning. Inspired by our earlier finding that benign taste experience increases the discriminability of gustatory cortical (GC) ensemble and single-unit responses (Flores et al., 2022), we hypothesized that taste experience could reshape the way that GC processes a taste stimulus by decreasing response variability. To test this hypothesis, we investigated how taste experience alters the GC taste response to a novel taste (i.e., sucrose) both before (pre) and after (post) CTA conditioning.

The previous report demonstrated that benign experience impacts the processing of both familiar and novel tastes (Flores et al., 2022); novel sucrose responses became more discriminable and distinct from the response towards water. The current Fano Factor analysis reveals that this enhancement in discriminability is associated with (and likely driven by) an increase in the reliability of taste coding. Specifically, trial-to-trial sucrose responses in GC pre-CTA are less variable in the taste-experienced group than in the taste-naïve group. These results imply that taste experience enhances the reliability of the GC response, increasing signal to noise ratios.

In addition to single unit activity, GC ensemble activity also becomes more stable following taste exposure. As noted previously, GC taste dynamics progress through distinct epochs, encompassing detection, identity, and palatability (Katz et al., 2001; Jones et al., 2007; Sadacca et al., 2016). The current work demonstrates that taste experience also impacts the across-neuron coherence of single-trial transitions into an epoch previously shown to be critical for CTA learning (Moran and Katz, 2014), increasing the across-ensemble coherence of this firing-rate change. Given that the cohesion of an ensemble’s activity can be linked to an enhanced probability of plasticity (Li, 2018), it is reasonable to suggest that taste experience boosts learning towards the conditioned stimulus (sucrose) by improving the quality (i.e., coherence) of epoch transitions in GC.

It has been well established that reduced variability (both in terms of stimuli and neural responses) and learning go hand-in-hand (see Raviv et al., 2022 for a review). For instance, exposure to multiple tastes at the same time of aversion conditioning towards novel sucrose creates an overshadowing event; the intended conditioned stimulus is overshadowed by the presence of the other tastes. Presenting multiple tastes simultaneously inherently enhances variability of neural activity (by driving multiple taste responses) leading a reduced aversion learning towards any of the presented tastes (Pavlov, 1927; McLaren and Mackintosh, 2002; Flores et al., 2016). When considering the natural response variability towards multiple trials of the same stimulus, it’s important to consider the possibility that even this type of variation might make learning sub-optimal as the variation may produce an overshadowing effect by rendering the system to treat each taste trial, even if the taste is the same, as slightly different stimuli (Shadlen and Newsome, 1994, 1995). Thus, the fact that water-experienced rats showed much higher variability within CS-evoked firing rates across trials than those that had taste experience before conditioning can explain a weaker learning towards novel sucrose compared to experienced rats.

Interestingly, in addition to altering evoked responses, benign taste experience also alters response variability during spontaneous activity (when the stimulus is not presented). Like the impact on taste responses, rats exposed to the taste battery also show reduced variability in baseline activity compared to rats naïve to tastes. This impact has been observed in other sensory systems (e.g., vision; Miller et al., 2022; Niraula et al., 2023). Niraula et al. (2023) reported that following inconsequential exposure to visual stimuli, the spontaneous activity of visual cortex becomes more similar to the response induced by the pre-exposed stimuli. In other words, spontaneous activity becomes less random and shifts to resemble the responses towards stimuli. This shift, importantly, is functionally relevant and has been linked to enhanced novelty detection (Haider et al., 2007; Scholvinck et al., 2012). This enhancement in novelty detection may also account for our earlier finding that GC sucrose response is more discriminative (from water) in taste-experienced rats than those without the experience (Flores et al., 2022). Furthermore, in taste processing, novelty is often studied in terms of neophobia (the fear of novel tastes (e.g., Lin et al., 2009; Wiaderkiewicz and Reilly, 2023)) which bears a close relationship to CTA learning; novelty boosts the associativity of stimuli (Reilly and Bornovalova, 2005; Lubow, 2009). Thus, the impact of taste experience on GC spontaneous activity reported here, while unexpected, may also play a role in in the experience-dependent enhancement of later CTA learning. Future studies should evaluate whether changes in spontaneous activity plays an independent role in the modulation of aversion learning or whether its role is more interdependent where changes in spontaneous activity influence palatability processing, which in turn, impacts learning.

The studies that show an enhancement of CTA learning following taste experience involve the same pre-exposed tastes (i.e., salty, sour, and water) followed by an aversion conditioning to novel sucrose. Thus, it is possible that the reported neural findings here are associated to this specific combination of tastes and do not generalize to other tastes being pre-exposed or conditioned during CTA learning. This is unlikely given our previous work showing that exposure to a single taste alone induces LE towards a novel taste. Furthermore, preliminary evidence showing that experience with sour and sweet tastes still enhances an aversion to a novel salty taste (data not shown). While future research is still needed to continue to examine the bounds of benign taste experience’s impact on learning using different tastes, this preliminary evidence suggests that the impact of taste experience on the reliability of taste responses is not taste specific.

We focus on how taste experience impacts novel taste processing in the brain before and after aversion conditioning. Here, we report how these benign experiences significantly influence GC activity pre- and post-aversion learning towards a novel taste. While the current experimental design cannot provide direct evidence that the experience-induced changes enhance future CTA learning, the results from our ancillary analysis suggest these changes could, at least partially, contribute to learning enhancement. If the reliability of taste responses plays a role in CTA learning, then we should expect to observe better CTA learning in rats showing lower Fano Factor (i.e., less variation) in their taste response regardless of whether the rats is taste experienced or taste naive before CTA learning. This is exactly what is revealed from this analysis; how well the rats learned a CTA is correlated with the Fano Factor (Figure 6), calculated from the taste response within the palatability-related epoch during CTA learning (pre-CTA).

**Figure 6.**
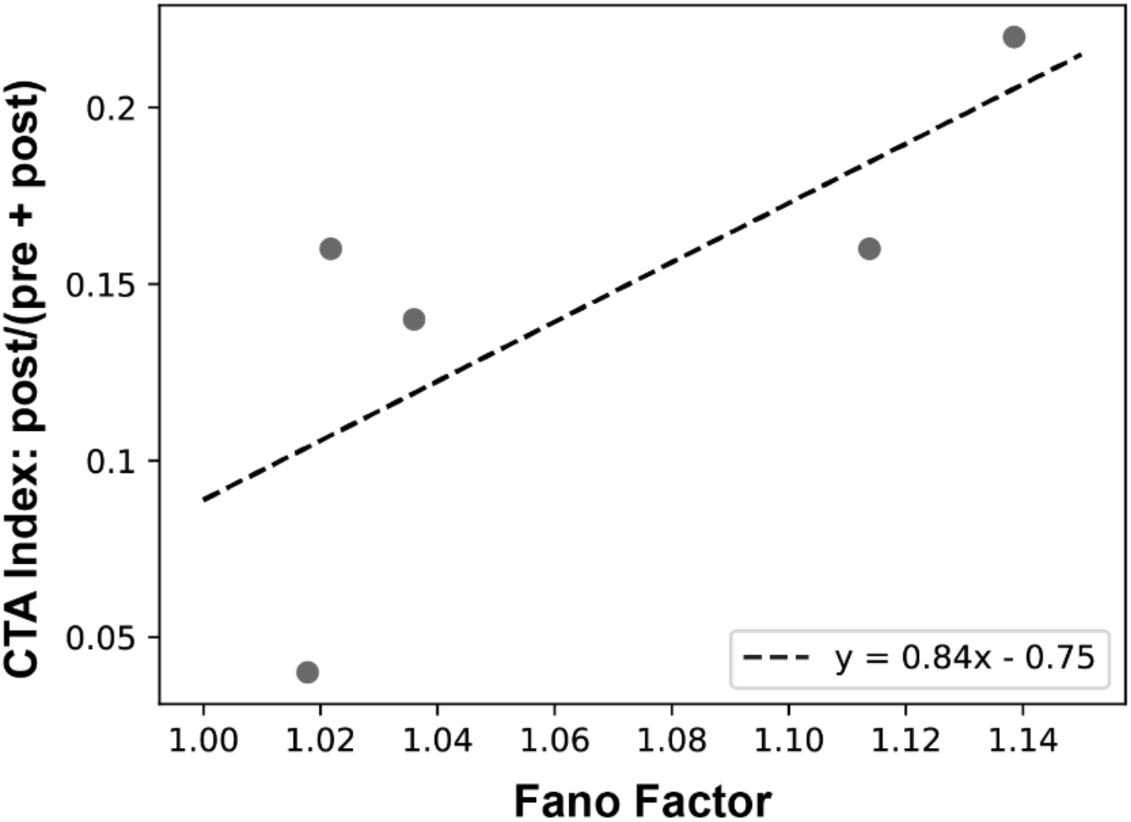
Strength of Aversion Correlates with Fano Factor Measure of Reliability. CTA strength correlates with the reliability of sucrose CS processing during CTA conditioning. For each rat for which behavioral data was available (indicated by each dot) a Fano factor (calculated from the firing rates during the palatability epoch [800-2000ms post-taste delivery] across trials) was plotted against CTA strength (lower CTA index = stronger CTA) for the same rat (*r* = 0.82, *p* = 0.044). There is a significant positive correlation between CTA index and FF values (the stronger the CTA, the lower the FF).

The finding that benign taste experience can influence how the brain processes a novel taste stimulus implies that such experience is not actually inconsequential. The impact of benign experience on sensory processing has also been reported in olfaction (Mandairon et al., 2006) and sensory development (Schiff et al., 2023), but remains understudied despite its omnipresence. Our current *in-vivo* electrophysiological study sheds light for the first time on potential neural mechanisms that might underlie these experiential and neural dynamics. Future research involving neuronal interventions (optogenetic or chemogenetic inhibition) will be required to further our understanding of the intricate neural circuitry and mechanisms responsible for the impact of “benign” experience on learning. This, in turn, may offer insights into how everyday experiences impact the neural dynamics of future sensory processing, consumption disorders, and learning and memory formation.

## Acknowledgement

We would like to sincerely thank Dr. Donald B Katz for his feedback on this manuscript. We gratefully acknowledge funding from the National Institute of Deafness and other Communication Disorder (NIDCD) through the National Institute of Health (NIH): Grant Numbers: R01 DC006666 and R01 DC007703 to DBK, R21 DC016706 to JYL, and F31 DC 015931 to VLF.

